# Dynamic Nature of *Staphylococcus aureus* Type I Signal Peptidases

**DOI:** 10.1101/2024.01.23.576923

**Authors:** Jesper J. Madsen, Wenqi Yu

## Abstract

Molecular dynamics simulations are used to interrogate the dynamic nature of *Staphylococcus aureus* Type I signal peptidases, SpsA and SpsB, including the impact of the P29S mutation of SpsB. Fluctuations and plasticity-rigidity characteristics vary among the proteins, particularly in the extracellular domain. Intriguingly, the P29S mutation, which influences susceptibility to arylomycin antibiotics, affect the mechanically coupled motions in SpsB. The integrity of the active site is crucial for catalytic competency, and variations in sampled structural conformations among the proteins are consistent with diverse peptidase capabilities. We also explored the intricate interactions between the proteins and the model *S. aureus* membrane. It was observed that certain membrane-inserted residues in the loop around residue 50 (50s) and C-terminal loops, beyond the transmembrane domain, give rise to direct interactions with lipids in the bilayer membrane. Our findings are discussed in the context of functional knowledge about these signal peptidases, offering additional understanding of dynamic aspects relevant to some cellular processes with potential implications for drug targeting strategies.

**Highlights:**

- Dynamics of *S. aureus* SpsA and SpsB (wild-type and P29S) is analyzed.
- Differences in flexibility and plasticity are relevant for function.
- P29S mutation decouples mechanical motions in SpsB.
- Extracellular domain loops insert differently into the membrane in SpsA and SpsB.
- *S. aureus* model membrane properties are elucidated.

## INTRODUCTION

*Staphylococcus aureus*, a Gram-positive bacterium, is responsible for causing various acute and chronic infections. This bacterium is known to secrete a number of virulence proteins to the host environment and the bacterial cell surface, which are essential for its pathogenicity.^1–3^ Proteins destined for the extracellular environment are synthesized in a precursor form with an N-terminally cleavable signal sequence, commonly referred to as the signal peptide.^4,5^ Type I signal peptidases, which can be considered unusual serine proteases carrying a Ser-Lys catalytic dyad, cleave off the signal peptide from the exported protein precursors, thereby producing the mature protein either during or shortly after export.^6,7^ Cleavage is typically required to release exported proteins from the membrane^8^, though there are other mechanisms such as excretion^9–11^.

SpsB is the canonical type I signal peptidase in *S. aureus* and essential for bacterial growth.^12^ *S. aureus* appears to have only one such functional peptidase (namely SpsB), whereas other bacteria species may have multiple.^13–15^ SpsB is responsible for processing the secreted precursor constructs by cleaving off the N-terminal signal sequence of both generic and YSIRK type, resulting in varied localization patterns.^16–21^. Inhibition of SpsB is considered a promising antibiotic strategy, as SpsB has a Ser-Lys catalytic dyad instead of the prototypical Ser-His-Asp triad found in eukaryotes. Moreover, SpsB contains a transmembrane domain and an extracellular enzymatic domain that is easily accessible to antibiotics. Arylomycins, a group of compounds produced by Streptomyces,^22,23^ represent the most promising class of SPase inhibitors.^24^ Unfortunately, despite their promise, arylomycins face limited efficacy against *S. aureus*.^25,26^ Compounding this challenge is the emergence of resistance mechanisms.^27,28^ However, certain naturally occurring variants P29S found in *Staphylococcus epidermidis* exhibit sensitivity to arylomycin.^29^ Introducing P29S mutation to *S. aureus* wild-type SpsB decreased resistance to arylomycins.^29,30^

*S. aureus* encodes another type I signal peptidase, SpsA. SpsA is a homolog of SpsB, which is located adjacent to *spsB* in staphylococcal genomes. SpsA is thought to be a non-functional or inactive type I signal peptidase due to the two conserved catalytic residues missing,^12^ and speculation is that is has no functional roles and may instead be the product of a pseudogene. Indeed, SpsA carries no peptidase activity in biochemical assays.^31^ Intriguingly, it was recently shown that SpsA contributes to *S. aureus* phagosomal escape,^32^ raising the question of its unknown function. It is likely that it may retain ability to bind various peptides or proteins in a way that is functionally relevant, only that it cannot catalyze the enzymatic peptidase reaction.

Structural analyses of *S. aureus* SpsB has shed light on similarities with other signal peptidases, such as *E. coli* LepB.^33–35^ These findings show that domain architecture as well as structural elements influencing peptide binding and arylomycin binding appear highly similar.^36^ However, limited attention had been given to elucidating the dynamic nature of the enzymes, which is the primary focus of the present investigation. For this purpose, we have conducted extensive molecular dynamics simulations on the two *S. aureus* signal peptidases SpsA and SpsB *apo*-enzymes, as well as the important P29S mutation.

## RESULTS AND DISCUSSION

### Structure and Dynamics

The structure of SpsB had been determined from X-ray crystallography, both in its unbound (apo) state and when complexed with specific signal peptide substrates, employing an elegant fusion approach.^34,35^ These structures have revealed a bipartite fold within the extracellular domain (ECD) of SpsB, which is homologous to *E. coli* LepB.^33^ However, notable differences exist in the noncatalytic segment of ECD, which may play a role in recognizing the mature protein component of the substrate.^37^ In addition, the catalytic dyad residues, Ser-36 and Lys-77 in SpsB, exhibit similar arrangements to those in LepB, forming the lining of the highly conserved substrate binding pocket.

The structure of the transmembrane domain (TMD) of SpsB is highly likely to consist of a single alpha-helical unit based on predictions from both secondary structure prediction algorithms and AlphaFold2. As a result, the TMD spans the entire cellular membrane, with a cluster of positively charged residues positioned N-terminally on the intracellular side. The ECD has limited lipid interactions deep within the membrane but for a couple of surface loops whose hydrophobic residues may insert peripherally into the membrane (Figure 1). A detailed exploration of these interactions is presented in a section below.

**Figure 1:**
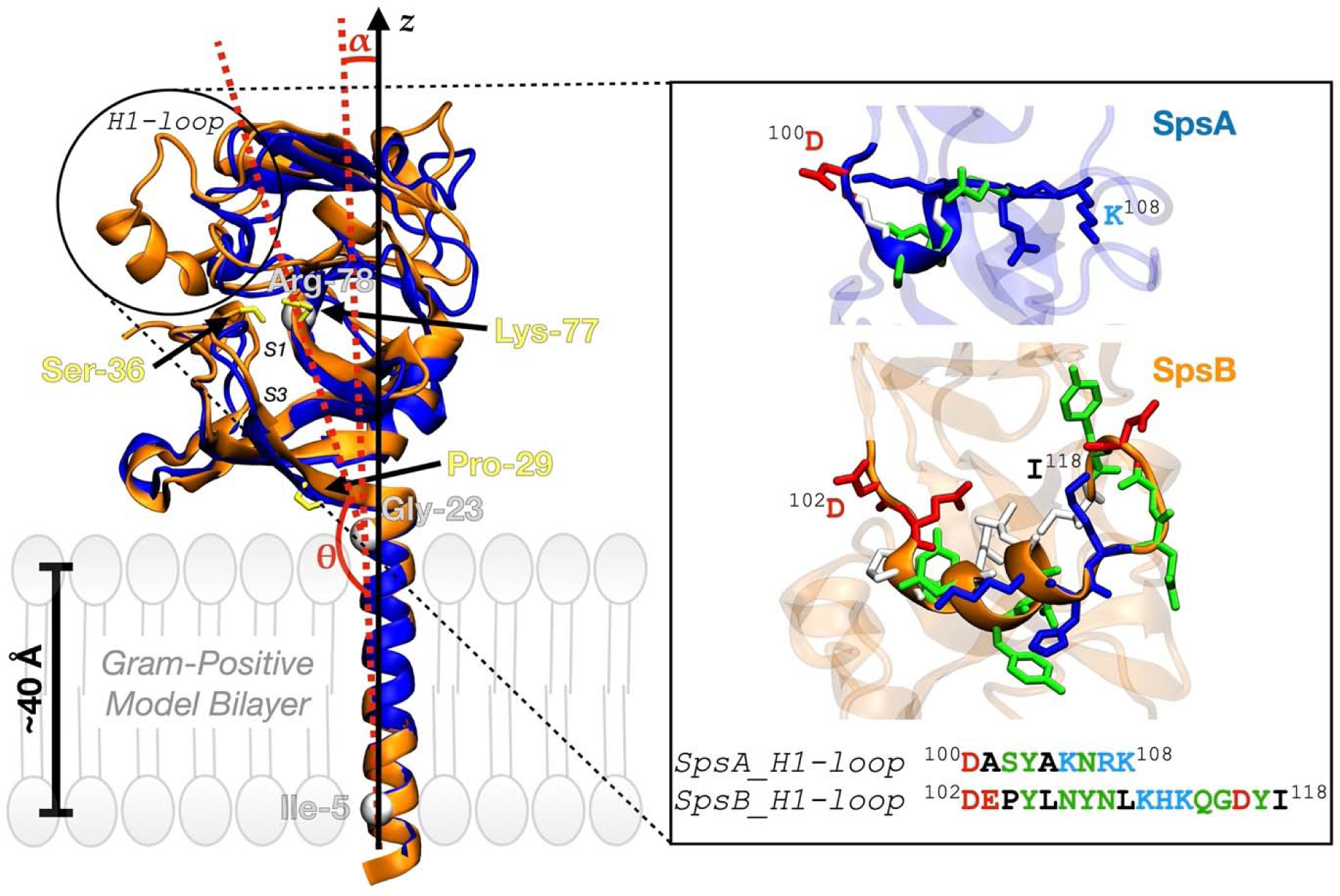
Schematic depiction of the initial setup of SPases (shown in cartoon representation with SpsA in blue and SpsB in orange) with respect to the membrane surface. Key residues are highlighted, including the active site residues (Ser-36 and Lys-77, here and forthwith numbering will correspond to SpsB unless otherwise specified) and the position of the arylomycin-sensitive mutation in SpsB (Pro-29), as well as alpha-Carbon atoms of Arg-78, Gly-23, and Ile-5 (as spheres, in gray) used for calculation of the TMD tilt angle, a, and the inter-domain bend angle, 0. The SPase is positioned such that the TMD traverses the membrane with a tilt angle of a = 0°. The bilayer thickness is ∼40 Å (see also Figure 6c). The zoomed-in region focuses on the helix 1 (H1)-loop region, with the corresponding structure and sequence for SpsA and SpsB shown in the right-side panel. Amino acid coloring in the panel reflects the chemical properties of residues (negatively charged in red, positively charged in blue, hydrophobic in white/black, polar in green).

SpsA is similar to SpsB in terms of both sequence (31% identity overall) and structural alignment (0.95Å RMSD over 139 alpha-Carbons). The major structural differences are found in helix 1 (H1) and the ensuing loop of the noncatalytic segment of the ECD (residues 102-118 in SpsB; residues 100-108 in SpsA), where SpsA features a truncated version as compared with SpsB (Figure 1).

The influence of the P29S mutation in SpsB affect arylomyin binding, making it sensitive to arylomycin antibiotics.^29^ However, it is not expected to have significant influence on the overall structure of the peptidase,^36^ apart from introducing a local perturbation due to the distinct preferences of Proline vs. Serine. Since Tyr-30 lines the S3 pocket in SpsB, it is tempting to speculate that this pocket may be affected by the P29S mutation in a manner that makes it susceptible to arylomycin through structural changes around S3. However, as we shall see, this scenario may be overly simplistic as there are substantial differences in dynamics between WT and SpsB_P29S, with seemingly SpsA sharing more similarities with P29S rather than WT SpsB in certain aspects that will be detailed below.

### Molecular flexibility and fluctuations

Our initial focus is on investigating the flexibility and fluctuations exhibited by the three proteins. These characteristics directly influence the conformational landscape of the proteins. Despite sharing a similar structural topology, there are notable distinctions between SpsA, SpsB, and SpsB_P29S. As previously mentioned, the difference (at the sequence level) between SpsA and SpsB is mainly in the H1-loop segment. In contrast, the difference between SpsB WT and SpsB_P29S is limited to a single point mutation. However, these differences manifest themselves in the molecular flexibility among the proteins individual loops have even been known to control enzyme activity and regulation.^38,39^ Although the structures overlap well when superimposing the transmembrane domain (TMD), there appears to be minimal difference in both TMD and apparently similar flexibility in the extracellular domain (ECD) (Figure 2a). However, aligning the ECD unveils that this similarity is due to the amalgamation of relative interdomain and intradomain fluctuations. When the former is reduced through structural alignment, crucial distinctions emerge in the H1-loop segment, exhibiting increased flexibility in SpsB. Meanwhile, SpsA and SpsB_P29S visually appear similar overall, including in the H1-loop segment (Figure 2a).

**Figure 2:**
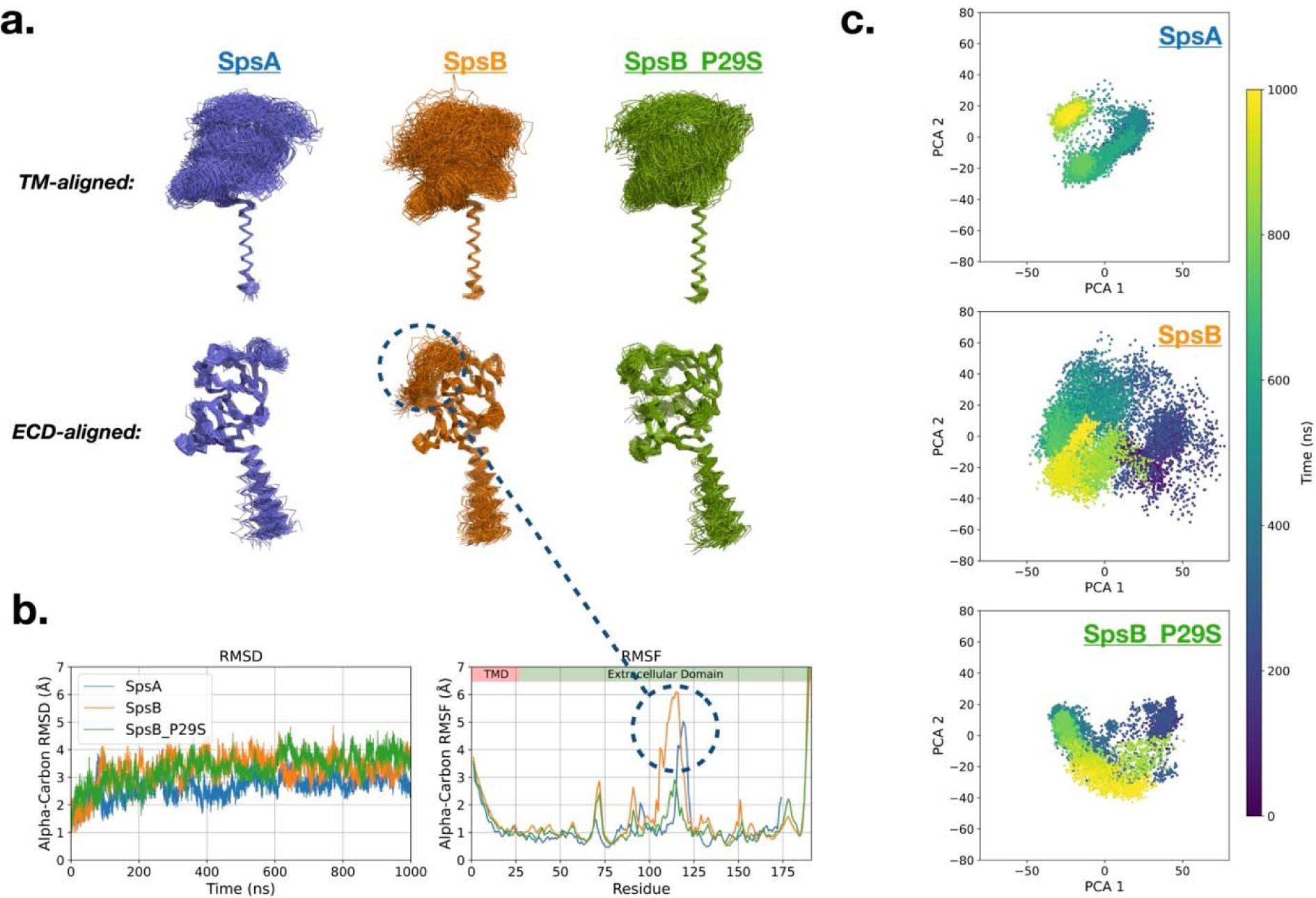
Dynamics of SPases. (**a**) Superimposed structural snapshots shown as ribbon diagram taken at 10-ns intervals from the simulations of SpsA (in blue), SpsB (in orange), and SpsB_P29S (in green) bound to the model S. aureus membrane. Superimposition was performed on the TMD for the top ensembles and on the ECD for the bottom ensembles. (**b**) Plots showing the structural deviation from initial structure (root mean squared deviation, RMSD) and the structural fluctuation for protein residues (root mean square fluctuation, RMSF). (**c**) Principal component analysis of the simulation trajectories (ECD only). The plots show the first (PC1) and second (PC2) principal component against each other, with the color bar scale indicating the simulation time from 0 ns (in purple) to the end of the trajectory (in yellow). Additional principal components (PC1-3) can be seen in Figures S1, S2, and S3 in the supporting materials for SpsA, SpsB and SpsB_P29S, respectively.

Based on this, we anticipate limited structural deviation compared to the starting structures, and this is indeed that we observe in the root-mean-square deviation (RMSD) plot, where the overall alpha-Carton RMSD stabilizes at around ∼3Å for all simulations (Figure 2b). Quantifying fluctuations on the per-residue level identified the specific segments in the three proteins that are more or less flexible. SpsA and SpsB_P29S are exhibit overall similarity, reaching RMSF values as high as ∼6Å (Figure 2b). In contract, the corresponding region in P29S peaks at RMSF values less than 3Å. Notably, the shortened H1-loop segment in SpsA results in a shift in the local maximal fluctuation to the subsequent loopy segment around residues ∼113-122. Otherwise, all other regions appear with similar of identical ranges of RMSF values among the three proteins.

### Plasticity-rigidity axis

Next, we investigated the concepts of plasticity and rigidity, which are important factors in understanding the structural and functional aspects of proteins.^40–43^ Though, the terms are ill-defined despite being commonly used. We adopt the definition proposed by Csermely, which posits that plasticity and rigidity are opposing characteristics of the system due to external or internal changes.^44^ Structural plasticity and rigidity, defined in terms of structural properties, may not necessarily align with their functional counterparts.^45–48^ To explore these dynamics, we performed principal component analysis (PCA) on the simulations of the three peptidases (Figure 2c). PCA is useful technique for reducing the dimensionality of MD simulation data, allowing us to discern larger amplitude motions and compare the dynamics in the essential subspace.^40,41^

The first few principal components (PCs) are expected to effectively capture major conformational changes between distinct states,^40,41^ while the dynamics of a rigid protein within a narrow energetic well may involve independent motions of various protein parts, resulting in higher dimensionality in the essential dynamics space and local frustration.^49,50^ The PCA results indicate that SpsA and, to a lesser extent, SpsB_P29S are characterized by isolated and compact clusters, suggesting a high degree of conformational rigidity. In contrast, SpsB exhibits a different pattern with widely distributed points captured by PC 1 and PC 2, indicative of predominant conformational plasticity. We are tempted to interpret this finding as suggesting that wild-type SpsB, with its plasticity, may be better equipped to adapt to a diverse array of binding partners, such as substrates, compared to SpsB_P29S and SpsA. Though, we are not aware of any definitive evidence regarding substrate specificity or the expected number of binding partners between wild-type and P29S variant of SpsB.

The observation for SpsA, on the other hand, aligns with the presumption that it is not a functional peptidase in terms of enzymatic activity. However, it remains mysterious whether it has other unknown functions. Notably, the *spsA* mutant showed drastically decreased phagosomal escape.^32^ Although SpsA does not catalyze enzymatic reactions, it likely engages with other protein partners in a functionally relevant manner.^32^ Extended results of the first three principal components for each peptidase are available for SpsA (Figure S1), SpsB (Figure S2), and SpsB_P29S (Figure S3), all consistent with the aforementioned observations.

### Integrity of the active site

The structural integrity of the active site is important in facilitating catalytic competency.^51^ There exists surprising variation in the configuration of active sites among proteins, with some necessitating the binding of cofactors to induce structural and dynamic changes in the active site or other enzymatic hallmark features (e.g., substrate specificity pockets or oxyanion hole) to mature the enzyme and prepare it for substrate binding and hydrolysis.^46,51,52^ In the case of SpsA, the absence of the Ser-Lys dyad residues in the active site leads us to explore the corresponding residues at those positions, which would be Asp-35 and Ser-75, to which we refer as the ‘active site’ residues for simplicity.

By comparing the computed distances between gamma atoms (gamma-Carbon or gamma-Oxygen in the case of Ser) during simulations (Figure 3a), distinctions emerge in the conformations in the three peptidases. Specifically, the local environment appears quite similar in both SpsB and SpsB_P29S, while SpsA displays markedly different conformations in the active site. Three main conformational populations are identified as “inactive”, with distances at around 7-8 Å, “active” with distances at around 5-6 Å, and “compacted” with distances at around ∼4 Å (Figure 3b). SpsA predominantly resides within the inactive basin, whereas SpsB and SpsB_P29S primarily occupy the active basin, with minor fluctuations exploring the other two basins (Figure 3a).

**Figure 3:**
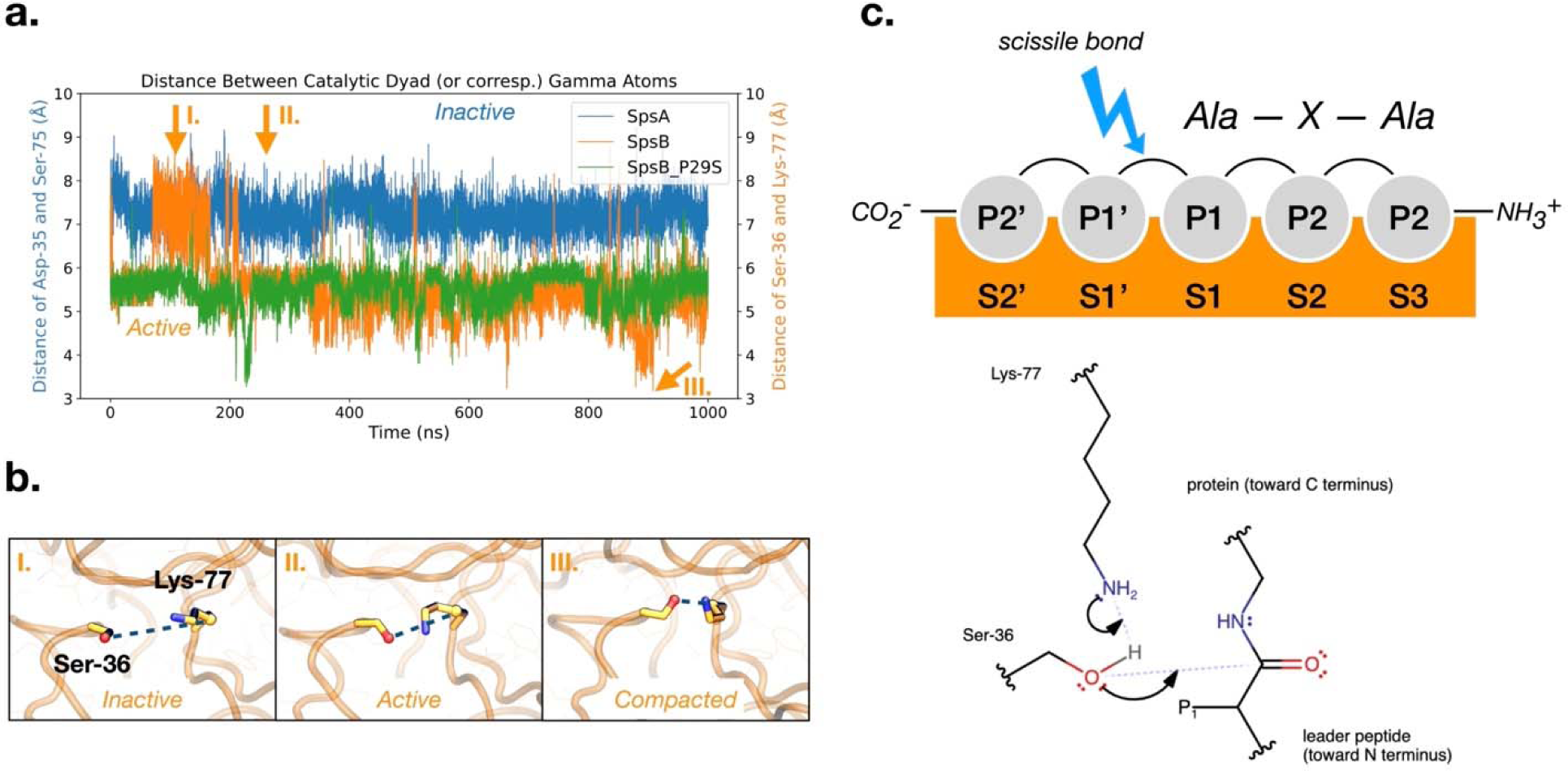
Configuration of the active site dyad residues and enzymatic formation of the Michaelis-Menten complex. (**a**) Time course plot showing the distance between gamma atoms of the two active site residues Ser-36 and Lys-77 (for SpsB, in orange, and SpsB_P29S in green), or Asp-33 and Ser-75 (for SpsA, in blue). (**b**) Structural snapshots of the active site conformations observed during the SpsB simulation, corresponding to “I” (inactive), “II” (active), and “III” (compacted) sampled basins. (**c**) Overview of the initial step in the enzymatic peptidase reaction, highlighting the scissile bond, substate specificity pockets on the primed and non-primed sides (S’ and S, respectively), and the nucleophilic attack by Ser-36 on the peptide’s carbonyl carbon and the of Lys-77 in enhancing Ser-36’s nucleophilic activity.

As noted earlier, the structural integrity of the active site dyad is essential for catalytic competency.^51^ The reason for this is that correct substrate alignment in space is required for the dyad residues to be coordinated to facilitate the catalytic reaction. In the peptidase reaction, the dyad residues must initially position themselves to form the Michaelis-Menten complex, a crucial step whose initial reaction mechanism involves the nucleophilic attach of Ser onto the carbonyl carbon of the substrate at the site of the scissile bond (Figure 3c). The role of Lys herein is to “activate” the Ser residue and enhance its nucleophilic properties (Figure 3c).

### Detailed membrane interaction and membrane-bound orientation

The *S. aureus* peptidase is a transmembrane protein, and the presence of the membrane is important in influencing both the structural configuration and dynamic behavior of the protein.^53,54^ In addition, the membrane environment may likely play a role in affecting the protein localization, molecular orientation, and functions through various mechanisms (e.g., by an allosteric effect).^53,55,56^ Indeed, it has been shown that phospholipids and the detergent Triton X-100 can increase the activity of truncated SpsB lacking TMD.^57,58^ We therefore explored the TMD interaction with the membrane, protein’s domain-domain bending in the membrane, known to have functional consequences for some proteins,^59,60^ as well as the detailed interaction between the ECD and the membrane.

### Mechanical coupling of TMD and ECD, and the influence of P29

The TMD domain consistently maintains an alpha-helix structure across all systems, functioning as a membrane anchor for the peptidase. The length of the TMD enables it to span the entire membrane, with positively charged residues located at the N-terminus exposed to the aquatic environment of the cytosol (Figure 1). However, because there are minor differences in the chemical composition of the two TMDs (SpsA and SpsB), the most favorable tilt angles are not precisely identical for the different proteins (Figure 4a).

**Figure 4:**
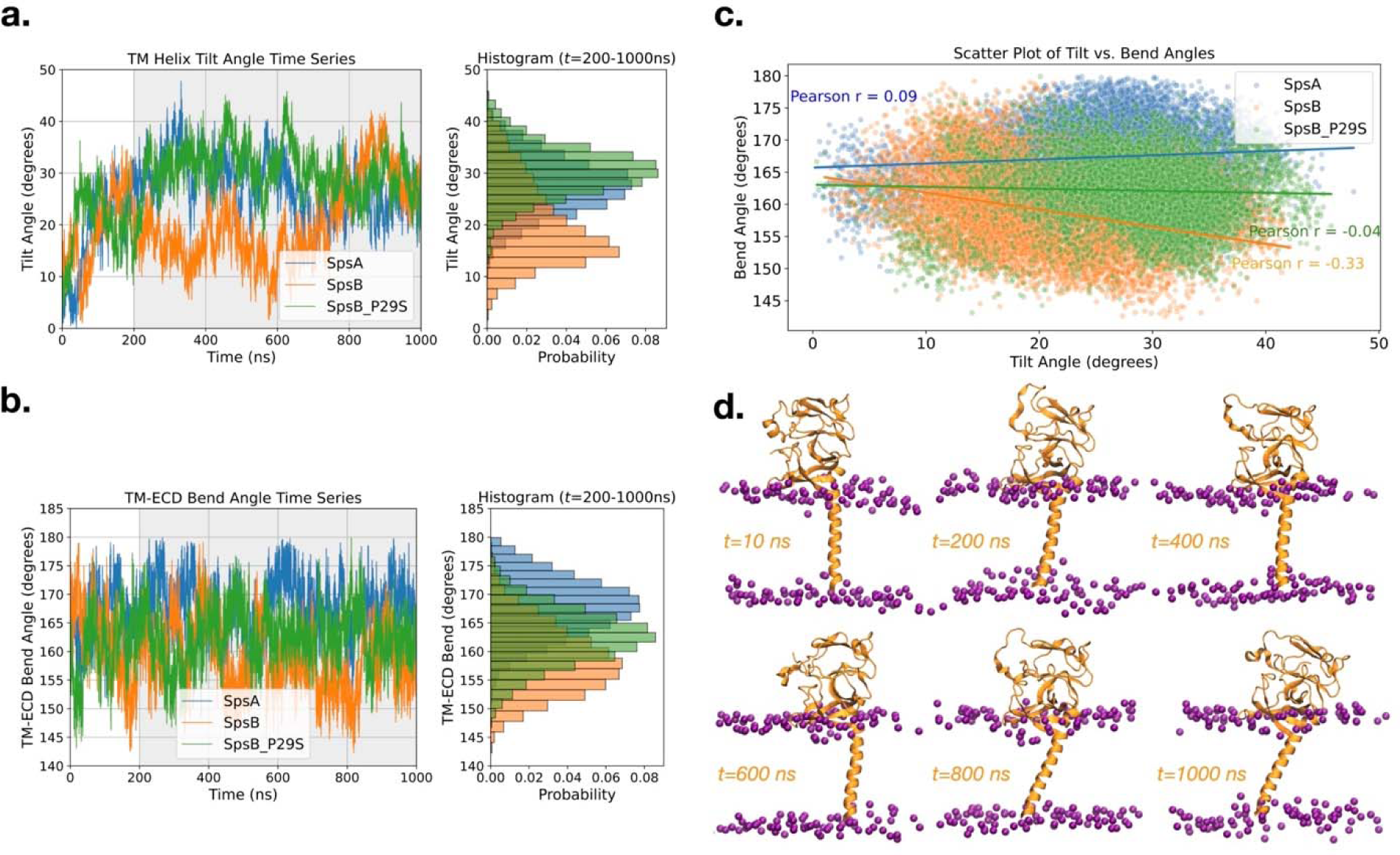
Molecular alignment of domains and their relative orientation to the membrane surface. (**a**) Time course plot of the TMD tilt angle, a, for the simulation of SpsA (in blue), SpsB (in orange), and SpsB_P29S (in green). The region shaded in gray is binned and shown in the righthand-side histogram. (**b**) Time course plot of the TMD-ECD inter-domain bend angle, 0, for the simulation of SpsA (in blue), SpsB (in orange), and SpsB_P29S (in green). The region shaded in gray is binned and shown in the righthand-side histogram. (**c**) Scatter plot illustrating the correlation between the tilt and bend angles, with the computed Pearson correlation coefficients. (**d**) Simulation snapshots of SpsB (shown as cartoon, in orange, along with the lipid headgroup P atoms shown as spheres, in purple) with corresponding timestamps.

There exists the possibility of a mechanistic coupling wherein the bending of protein domains, observed to vary during simulations (Figure 4b), may be interconnected and change in tandem with the TMD tilt angle. Our analysis reveals a weak correlation between the tilt angle of the TMD and the interdomain bend motion in SpsB (Figure 4c). The correlation has a Pearson correlation coefficient (R) of −0.33, suggesting a weak but discernible correlation. For SpsA and SpsB_P29S, these two properties are essentially uncorrelated (with Pearson’s R values of 0.09 and −0.04, respectively).

We speculate that the P29S mutation releases this mechanistic coupling, perhaps due to the additional allowed backbone configurations conferred by Ser compared to Pro, and the proximity of P29 to the connection between TMD and ECD. This notion aligns with the results for SpsA, which lacks a Pro residue at the corresponding position. Structural snapshots from the simulation of SpsB display the variability sampled and the synchronicity in domain-domain bending and TMD tilt (Figure 4d).

### Membrane-inserted residues at the 50s loop and the C-terminal loop

To further elaborate on the protein-membrane interactions, we interrogate the protein segments that are proximal to the membrane. This involves computing the relative height of each amino acid residue along the protein sequence, measured by the Z coordinate (with Z=0 centered in the middle of the membrane; see Figure 1). The results reveal that, in addition to the TMD, two segments of the proteins interact directly with the membrane lipids (Figure 5a). These are the 50s loop, connecting beta-sheets 2 and 3 of the catalytic part of the ECD, and the C-terminal loop of the ECD. Sequence variations give rise to distinct chemical footprints of these loops that affect the chemical details of the interactions with the membrane environments in general and specific lipids.

**Figure 5:**
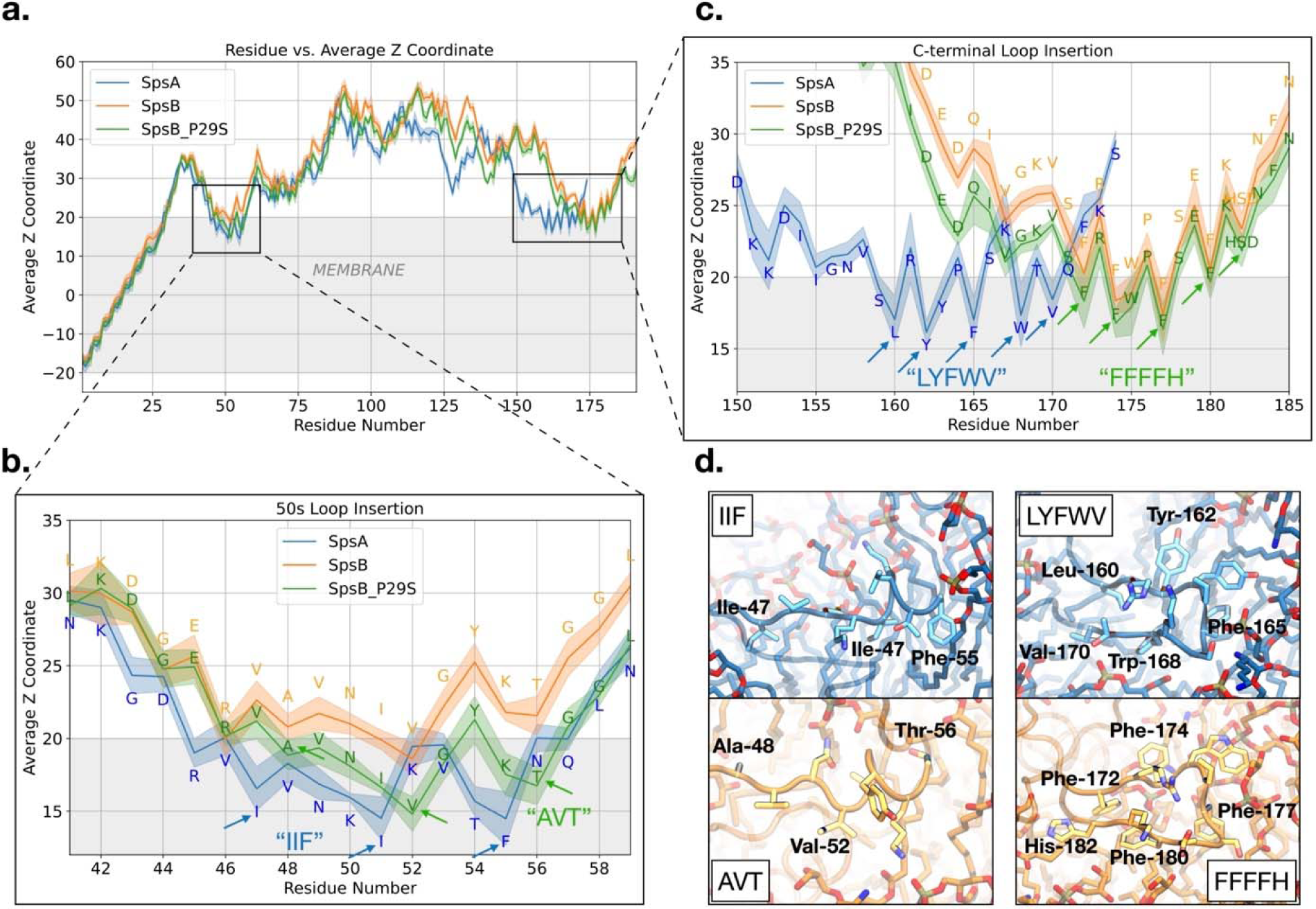
Elaboration of membrane-interacting residues in SPases. (**a**) Plot showing the average Z coordinate of each protein residue for SpsA (in blue), SpsB (in orange), and SpsB_P29S (in green). The color-shaded regions around each curve indicate the standard deviation and the gray-shaded portion of the plot represents the membrane. Zoomed-in inserts corresponding to the region around the 50s loops (in panel **b**) and the C-terminal loop (in panel **c**) are also presented for clarity. (**d**) Structural snapshots taken from the simulations of SpsA and SpsB show the distinctive interaction between chemically diverse loops motifs and membrane lipids (“IIF” and “LYFWV” for SpsA’s 50s and C-terminal loops, respectively; “AVT” and “FFFFH” for SpsB’s 50s and C-terminal loops, respectively). Coloring in the sub-panels are consistent with the other plots (SpsA in blue and SpsB in orange).

Examining the 50s loop, a motif of Ile-Ile-Phe (“IIF”) is observed in SpsA, whereas the corresponding motif in SpsB is Ala-Val-Thr (“AVT”) (Figure 5b). Comparing these, “IIF” appears more hydrophobic, likely explaining why this segment in SpsA inserts more deeply into the membrane compared to SpsB_P29S. To explain why SpsB inserts shallower than SpsB_P29S, we speculate that the constrained inter-domain movements may influence the insertion depth in SpsB. The membrane insertion of the C-terminal loop segment remains largely consistent among the three peptidases, despite sequence variations. Here, also WT and P29S SpsB demonstrate similar behavior with comparable loop insertions. The characteristic insertion motif of SpsA is Lys-Tyr-Phe-Trp-Val (“LYFWV”), with its corresponding motif in SpsBs is Phe-Phe-Phe-Phe-His (“FFFFH”) (Figure 5c). Structural snapshots illustrating the contacts made by the motif residues in the 50s and C-terminal loops of the contacts made by the motif residues in the 50s and the C-terminal loops are shown in Figure 5d.

### Properties of the S. aureus model membrane

Model membrane compositions specific to the *S. aureus* membrane are not widely studied in the scientific literature. Therefore, we have provided a detailed characterization of the mixture employed in the current study (Figure 6a). An inherent challenge in such simple mixtures is the potential for undesirable phase behavior (e.g., phase transition from liquid-disordered to liquid-ordered or the gel) in the membrane, even when the primary components of the biological membrane are reasonably represented.^61,62^ This may happen because the absence of active mechanisms found in living organisms that maintain membrane integrity.^61,62^ So, when using model mixtures, we need to carefully ensure that the membrane remains in its fluid phase. To this end, we found that we could not use majority myristoyl chains (14:0), exactly because the corresponding bilayer membrane undergoes phase transition into a gel phase, as evidenced by the decrease in APL from is 65.1 Å +/− 1.00 Å for composition 1 to 52.9 Å +/− 0.840 Å for composition 2 (Figure 6b and Figure S4 in the supporting materials). This transition is also evident in the component histogram along the membrane normal (Figure 6c) and the computed nematic order parameters of the bilayer’s main component (Figure 6d). In contrast, a previous MD study of a model *S. aureus* membrane composition consisting of 54% PG, 36% lysyl-PG, and 10% cardiolipin with anteiso-myristoyl chains (14:0, with a methyl branch on the second-last tail carbon) did not appear to undergo a comparable phase transition.^63^ It is conceivable that factors such as a slightly elevated temperature (313 K compared with 303.15 K), minor compositional changes, and/or the methyl branching are responsible.

**Figure 6:**
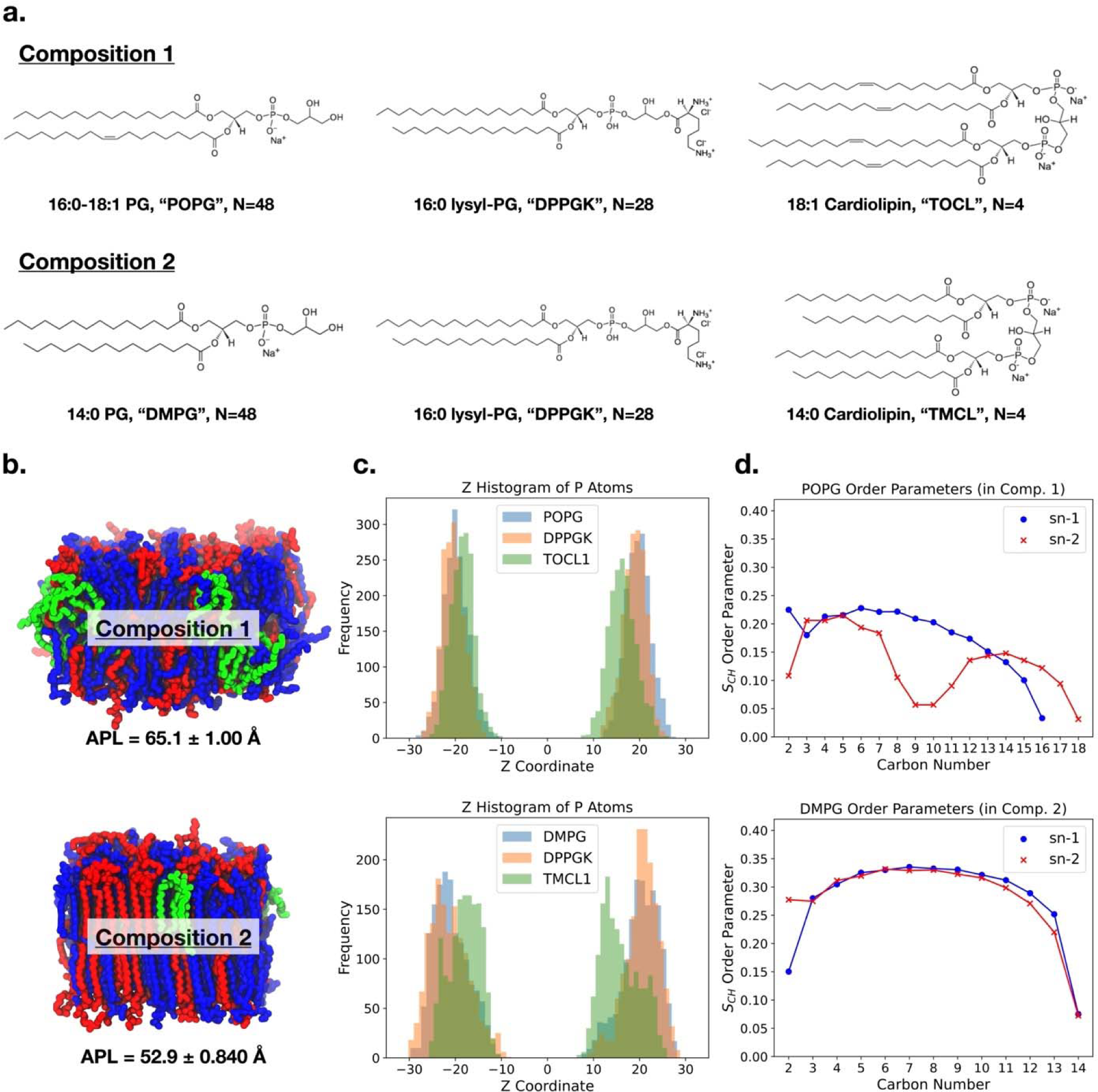
Evaluation of S. aureus model membrane compositions 1 and 2. (**a**) Presentation of the chemical structure and compositional details of the explored membrane compositions. (**b**) Structural snapshot of an equilibrated bilayer patch (after 500 ns) for composition 1 (top) and 2 (bottom), with membrane lipids shown as sticks (PG in blue, lysyl-PG in orange, CL in green). The respective equilibrated area-per-lipid (APL) is indicated below each snapshot. (**c**) Histograms showing the Z positions of lipid headgroup P atoms for the membrane compositions. (**d**) Analysis of unsigned lipid tail order parameters, S_CH_, for the *sn-1* and *sn-2* chains of the primary lipid constituent of the two compositions (POPG in comp. 1 and DMPG in comp. 2, respectively).

## CONCLUSION

In summary, our MD simulation results provide new insights into the dynamic nature behavior of the *S. aureus* type I signal peptidases SpsA and SpsB. Variations in fluctuations, plasticity-rigidity characteristics, and active site conformations underscore the diverse enzymatic capabilities of these peptidases. The observed alterations in mechanically coupled motions in SpsB, due to the P29S mutation, contribute to understanding of functionally relevant characteristics.

## METHODS

### Preparation of structures

SpsA and SpsB from *S. aureus* (strain Newman) were modeled as follows. The template for SpsB was derived from the crystal structure of the apo form of the peptidase (PDB ID: 4WVG).^35^ The missing segments in SpsB, as well as the entire structure of SpsA, were obtained from predicted structures available for the corresponding Newman strain UniProtKB entries (A0A0H3K7S6 for SpsA and A0A0H3KFC9 for SpsB) for the wild-type proteins (https://alphafold.ebi.ac.uk).^64,65^ The P29S mutation was introduced to SpsB *in silico*.

### Model membrane bilayer

The 60:35:5 ratio is a widely used ratio for components PG, lysyl-PG, and cardiolipin aimed at reproducing the properties of the *S. aureus* membrane in biochemical and biophysical assays.^66^ Maintaining the ratio, two different lipid mixtures differing solely in their lipid tail lengths (one being myriostol (14:0) acyl chains for PG and CL; the other having PO (16:0, 18:1) for PG and P (18:1) for CL; in both cases lysyl-PG was modeled as lysyl-DPPG (DPPGK, 16:0)). The resulting characterization of the *S. aureus* model bilayer is performed using in-house scripts as well as the NMRlipids Databank^67^.

### Molecular dynamics simulations

The MD box consisted of a model membrane patch, the protein construct, and saline TIP3P water^68^ with NaCl added to a physiological concentration of 150 mM. The transmembrane protein was placed in the membrane correctly by CHARMM-GUI builder^69,70^ and insertion depth determined by OPM^71^ with the TMD aligned with the *z*-axis. Forces were described using the CHARMM36m^72^ force field and production MD runs were performed in the NPT ensemble with pressure P = 1 atm and T = 303.15 K maintained by a Monte Carlo barostat^73^ and the Langevin thermostat^74^, respectively. The standard CHARMM-GUI protocol for minimization and relaxation were employed before initiating production run of 1,000 ns. All simulations were performed by OpenMM 7.5.3^75^ using particle mesh Ewald with a grid spacing of ∼1 Å for long-range electrostatics^76^ and switching off over the region from 10 Å to 12 Å. Periodic boundary conditions were applied in *x*, *y*, and *z* directions.

### Analysis of trajectories

Analysis of trajectories were conducted using VMD^77^ and in-house scripts relying on packages MDAnalysis,^78^ NumPy,^79^ pandas,^80^ Matplotlib^81^, seaborn^82^ and SciPy^45^. In addition, PyMOL (version 2.5.0, Schrödinger LLC) was used for inspection and visualization of proteins and membrane structures. Chemical diagrams were generated using Marvin (version JS 23.14.0, Chemaxon, https://www.chemaxon.com).

Tilt and bend angles were calculated based on position of alpha-Carbon atoms of Ile-5, Gly-23, and Arg-78 (see also Figure 1). The TMD tilt, a, is defined as the angle between the vector from Ile-5 to Gly-23 and the *z*-axis. The inter-domain bend angle, 0, is defined as the angle spanned by Ile-5, Gly-23, and Arg-78. The H1-loop is defined as the region from Asp-100 to Lys-108 and Asp-102 to Ile-118 in SpsA and SpsB, respectively. Principal component analyses of the construct ECD were computed over the backbone heavy atoms, starting at position 27 and the superimposed trajectory for that selection. The unsigned lipid tail order parameters for the *sn-1* and *sn-2* chains of the primary lipid constituent of the two compositions were computed as S_CH_ = (3 cos^2^0-1)/2, with angular brackets indicating molecular and temporal ensemble averaging.

### Authorship contribution

J.J.M.: Conceptualization, Methodology, Formal Analysis, Investigation, Visualization, Writing – Original Draft, Writing - Review & Editing W.Y.: Conceptualization, Writing – Original Draft, Writing - Review & Editing

### Declaration of competing interests

The authors declare no competing interests.

### Data availability

The simulation of the *S. aureus* model membrane is available at Zenodo (https://doi.org/10.5281/zenodo.8415046). Additional data that support this study are available from the corresponding authors upon reasonable request.

## Acknowledgements

All simulations are performed using the Advanced Computing Resources at University of South Florida, maintained by Research Computing. W.Y. is supported by the National Institute of General Medical Sciences (NIGMS-R35GM146993).

## Supplemental Information

**Figure S1:**
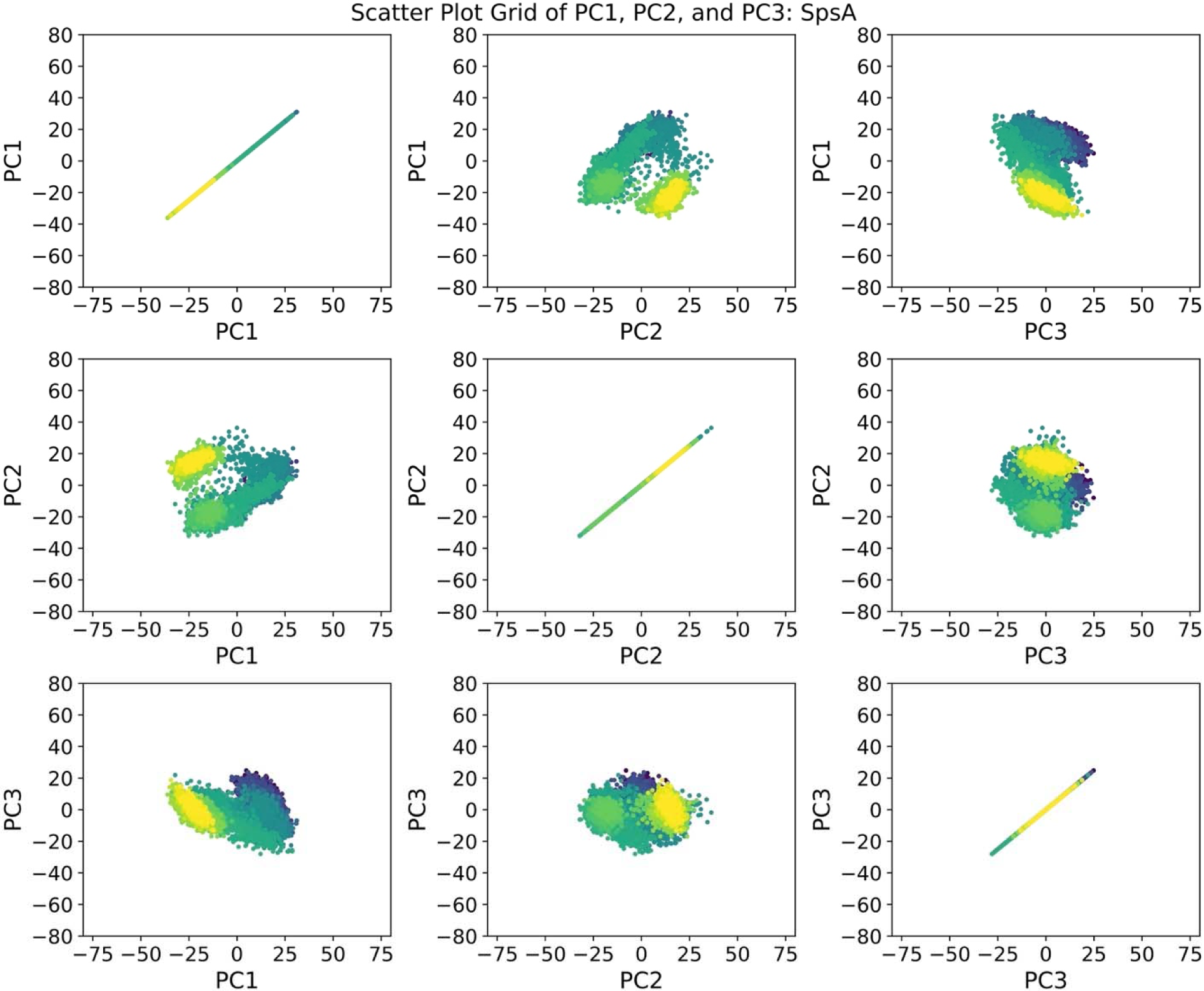
Extended principal component analysis showing a combinatorial scatter plot grid of PC1-3 for SpsA-ECD.

**Figure S2:**
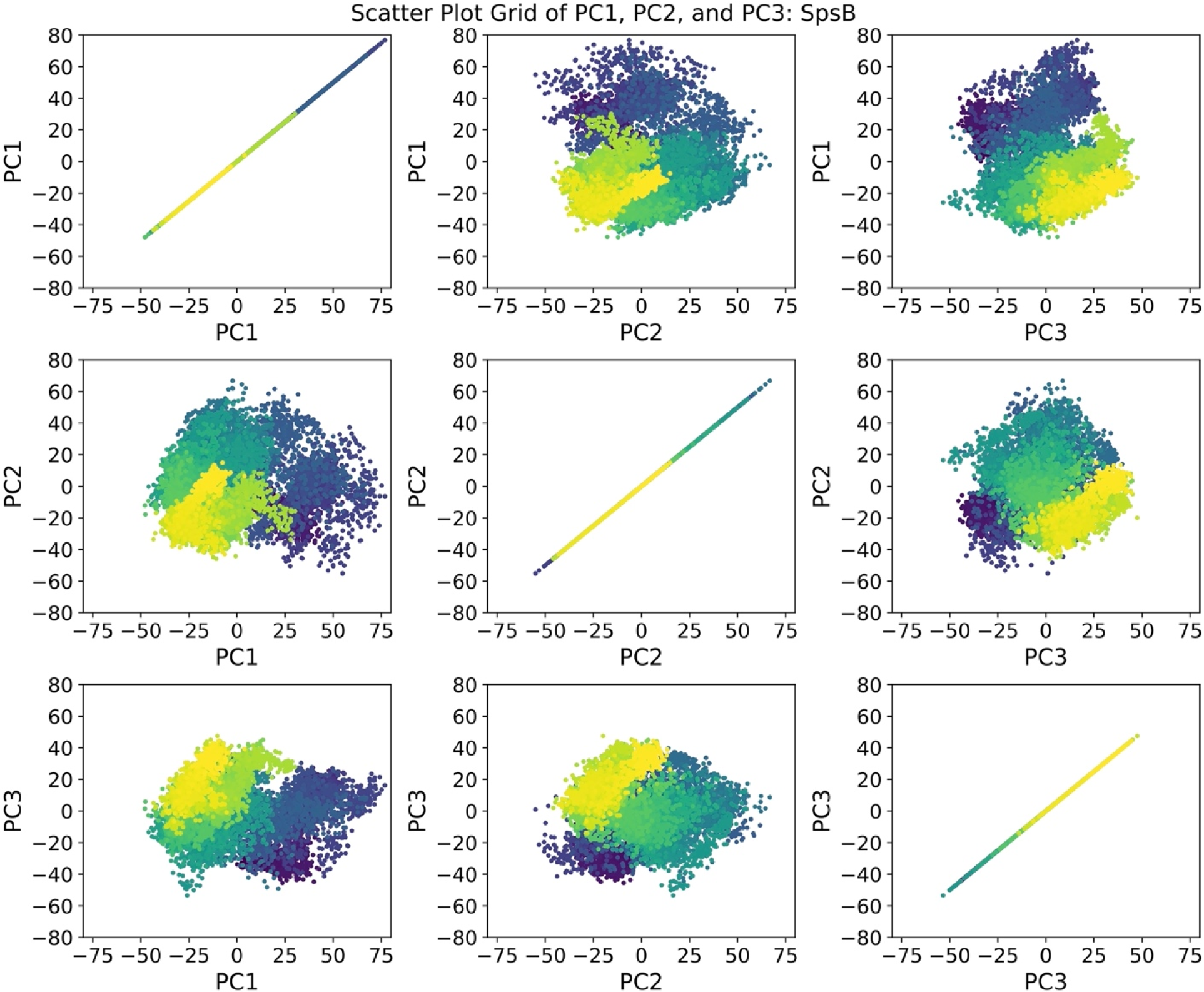
Extended principal component analysis showing a combinatorial scatter plot grid of PC1-3 for SpsB-ECD.

**Figure S3:**
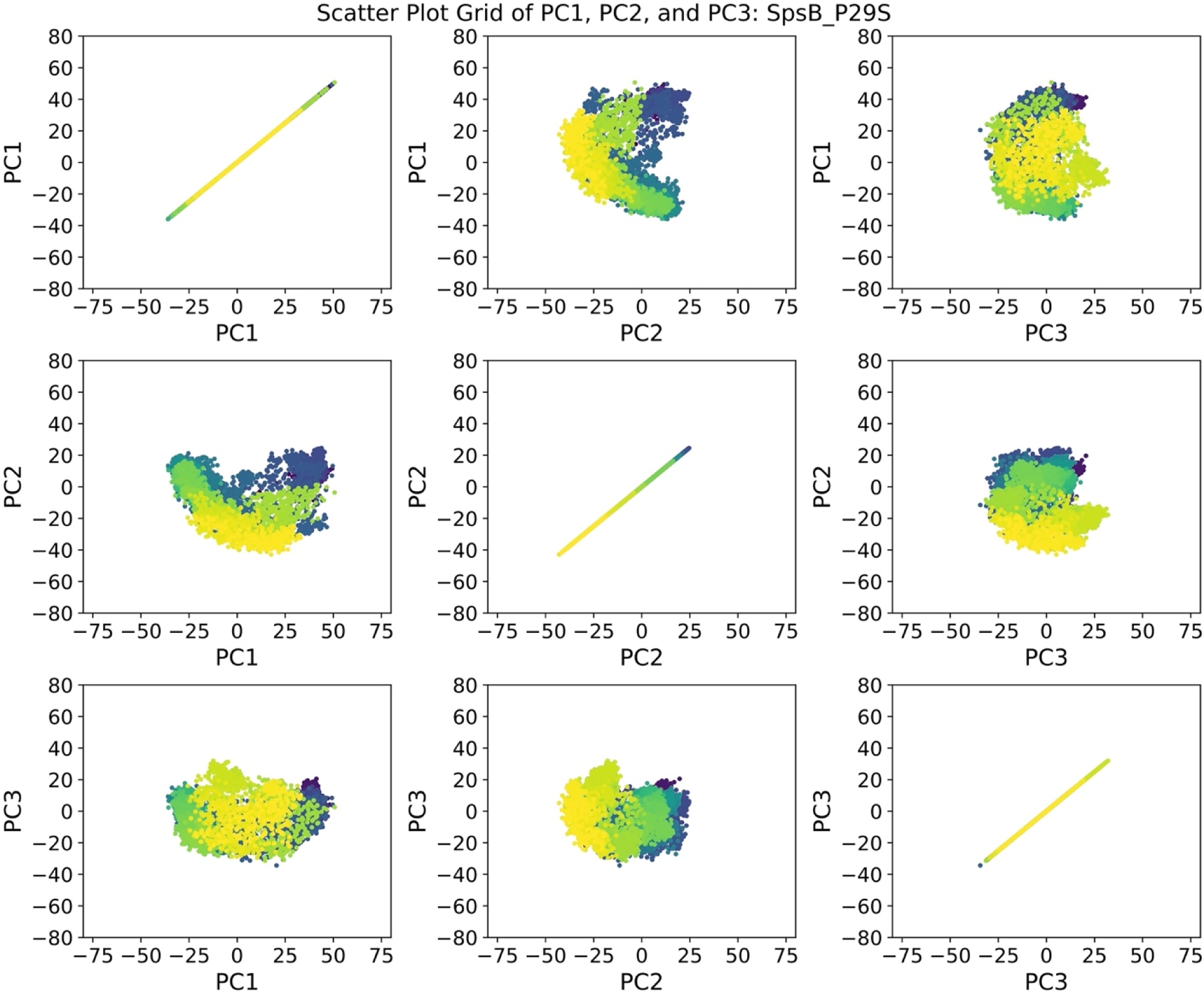
Extended principal component analysis showing a combinatorial scatter plot grid of PC1-3 for SpsB_P29S-ECD.

**Figure S4:**
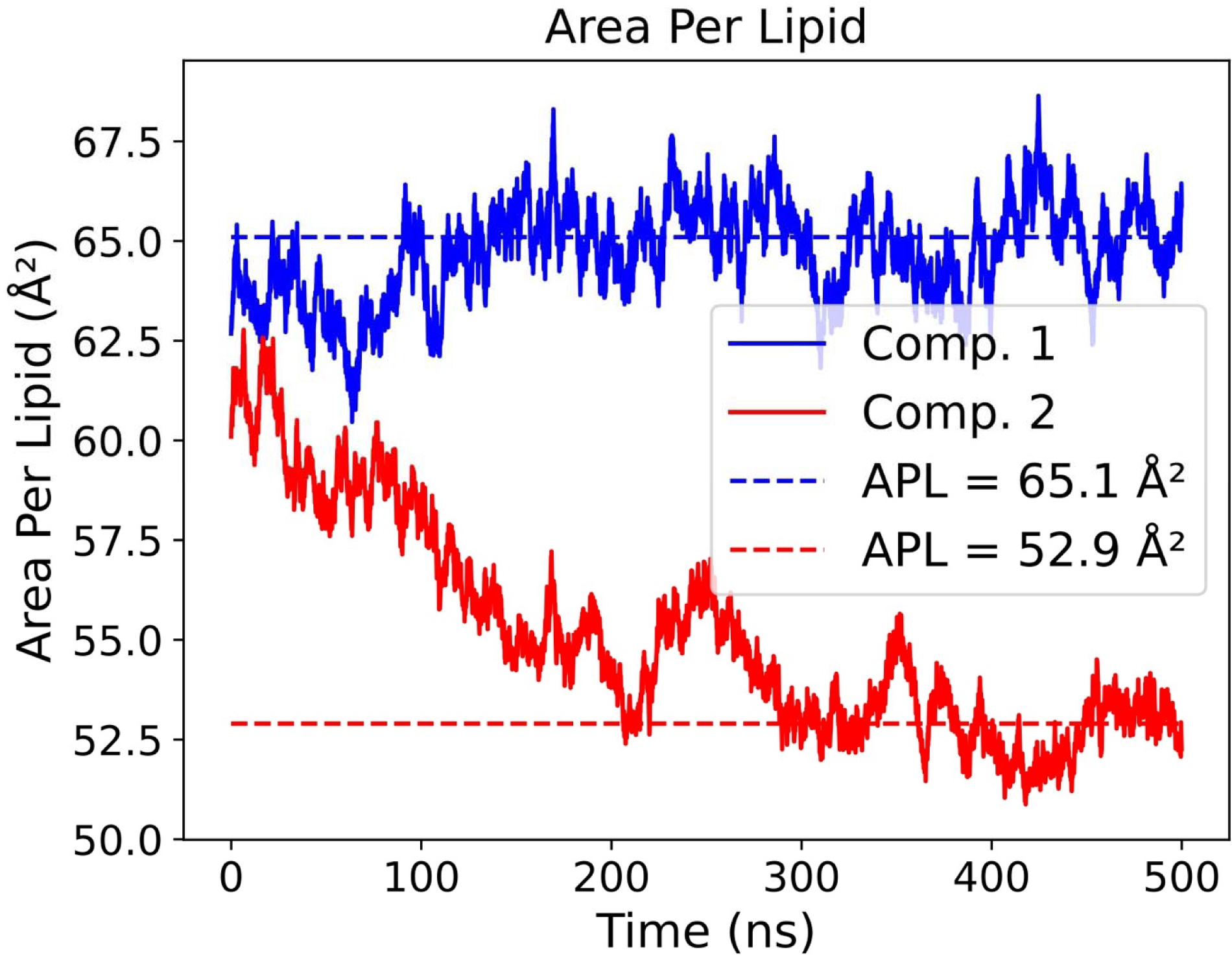
Area-per-lipid (APL) of membrane compositions 1 and 2 for a 500-ns equilibration MD run (without proteins present)

## Notes

### Competing Interest Statement

The authors have declared no competing interest.

